# Hippocampal disinhibition reduces contextual and elemental fear conditioning while sparing the acquisition of latent inhibition

**DOI:** 10.1101/2021.06.02.446736

**Authors:** Stuart A. Williams, Miriam Gwilt, Rebecca Hock, Charlotte Taylor, Joanna Loayza, Carl W. Stevenson, Helen J. Cassaday, Tobias Bast

## Abstract

Hippocampal neural disinhibition, i.e. reduced GABAergic inhibition, is a key feature of schizophrenia pathophysiology. The hippocampus is an important part of the neural circuitry that controls fear conditioning and can also modulate prefrontal and striatal mechanisms, including dopamine signalling, which play a role in salience modulation. Therefore, hippocampal neural disinhibition may contribute to impairments in fear conditioning and salience modulation reported in schizophrenia. To test this hypothesis, we examined the effect of ventral hippocampus (VH) disinhibition in male rats on fear conditioning and salience modulation, as reflected by latent inhibition (LI), in a conditioned emotional response procedure (CER). A flashing light was used as the conditioned stimulus (CS) and conditioned suppression was used to index conditioned fear. In Experiment 1, VH disinhibition via infusion of the GABA-A receptor antagonist picrotoxin prior to CS pre-exposure and conditioning markedly reduced fear conditioning to both the CS and context; LI was evident in saline-infused controls, but could not be detected in picrotoxin-infused rats due to the low level of fear conditioning to the CS. In Experiment 2, VH picrotoxin infusions prior to CS pre-exposure only did not affect the acquisition of fear conditioning or LI. Together, these findings indicate that VH neural disinhibition disrupts contextual and elemental fear conditioning, without affecting the acquisition of LI. The disruption of fear conditioning resembles aversive conditioning deficits reported in schizophrenia and may reflect disruption of neural processing within the hippocampus and its projection sites.

**Significance Statement:** Hippocampal disinhibition, reduced GABAergic inhibition, is a feature of schizophrenia, but how this contributes to psychological deficits remains to be clarified. Here, we focused on impairments patients show on classical-conditioning assays: aberrant salience allocation to stimuli that healthy participants have learnt to ignore and reduced fear conditioning, which have been linked to psychosis and negative symptoms, respectively. These impairments may be related to hippocampal disinhibition because the hippocampus modulates neural substrates of salience allocation and is part of the fear-conditioning neural circuit. Combining selective pharmacological manipulation of the hippocampus with a conditioning assay in rats, we found hippocampal disinhibition disrupted fear conditioning, without evidence for aberrant salience allocation. This suggests hippocampal disinhibition contributes to fear conditioning deficits in schizophrenia.

## INTRODUCTION

Hippocampal hyperactivity and neural disinhibition, i.e. reduced GABAergic inhibition, are key characteristics of schizophrenia pathophysiology and have been implicated in behavioural deficits characterising the disorder (Friston et al., 1992; Heckers & Konradi, 2015; Lieberman et al., 2018; Lisman et al., 2008; Tamminga et al., 2010; Tregellas et al., 2014). This hyperactivity is most evident in the anterior hippocampus (McHugo et al., 2019), corresponding to the rodent ventral hippocampus (VH) (Strange et al., 2014). Hippocampal disinhibition might contribute to behavioural impairments by disrupting neural processing both within the hippocampus, where regional disinhibition causes aberrant burst firing (McGarrity et al., 2017) and alters oscillatory activity (Gwilt et al., 2020) in rats, and in hippocampal projection sites (Bast et al., 2017; Katzel et al., 2020; Lodge & Grace, 2011). Here, we tested if hippocampal disinhibition contributes to deficits in latent inhibition (LI) and fear conditioning, which have been reported in schizophrenia.

LI refers to the reduced conditioning to a conditioned stimulus (CS), to which participants had been pre-exposed without consequence, and LI deficits have been reported in acute schizophrenia (Baruch et al., 1988; Gray et al., 1995; Rascle et al., 2001). One interpretation of reduced LI is that this reflects aberrant salience allocation to a stimulus that healthy participants had learned to ignore, and these findings contributed to the view that aberrant salience allocation is a key feature of schizophrenia and underlies psychotic symptoms (Gray et al., 1991; Howes et al., 2020; Kapur, 2003). Additionally, patients with schizophrenia show reduced aversive conditioning (Holt et al., 2009; Jensen et al., 2008; Romaniuk et al., 2010), which has been associated with negative symptoms (Holt et al., 2012).

The neural processes that underlie deficits in LI and aversive conditioning can be studied using rodent models. Permanent lesion studies in rats indicated that the hippocampus is not required for LI, although the adjacent entorhinal cortex and fibres passing through the hippocampus do play a role (Weiner, 2003); moreover, temporary inactivation studies indicated that the ventral subiculum may normally contribute to LI formation during pre-exposure (Peterschmitt et al., 2005; Peterschmitt et al., 2008). Interestingly, although NMDA-induced VH lesions spared LI acquisition, VH stimulation by local NMDA infusion moderately attenuated LI. However, this could partly have reflected reduced aversive conditioning (Pouzet et al., 2004). Although processing within the hippocampus could play a limited role in LI, VH stimulation and neural disinhibition might disrupt LI by stimulating dopamine release in ventral striatum and medial prefrontal cortex (mPFC) (Bast, 2011; Floresco et al., 2001; Legault et al., 2000; Mitchell et al., 2000; Peleg-Raibstein et al., 2005). Increased dopamine function, especially in the ventral striatum (Joseph et al., 2000; Nelson et al., 2011; Young et al., 2005), but also the mPFC (Morrens et al., 2020), has been shown to disrupt LI at conditioning. Additionally, VH disinhibition disrupted mPFC-dependent attention, presumably by way of strong hippocampo-mPFC projections (McGarrity et al., 2017; Tan et al., 2018), and could also disrupt LI acquisition during CS pre-exposure, which has been shown to require the mPFC (Lingawi et al., 2018). Apart from LI, VH disinhibition may also disrupt aversive conditioning itself, because the VH contributes to fear conditioning (Bannerman et al., 2004; Fanselow & Dong, 2010) and VH stimulation by NMDA was found to disrupt fear conditioning (Zhang et al., 2001).

Therefore, we tested the hypothesis that VH disinhibition would disrupt the acquisition of LI and fear conditioning in rats. We determined the effect of VH neural disinhibition via local microinfusion of the GABA-A receptor antagonist picrotoxin (McGarrity et al., 2017) on LI and fear conditioning, using a conditioned emotional response (CER) procedure with a CS pre-exposure stage (Nelson et al., 2011). Experiment 1 examined VH disinhibition during both pre-exposure and conditioning; this markedly reduced fear conditioning to the CS, so we were unable to examine changes in LI. Therefore, experiment 2 examined the effect of hippocampal disinhibition during pre-exposure only on the formation of LI.

## METHODS

### Rats

Overall, we used 104 male Lister Hooded rats (Charles River, UK), weighing 310-400g (9-12 weeks old) at the start of experiments. In Experiment 1, 72 rats were tested in 3 batches of 24 rats. Experiment 2 used 32 rats in a single batch. See the section on Experimental Design for further detail and for sample size justifications.

Rats were housed in groups of four in individually ventilated “double decker” cages (462 mm x 403 mm x 404 mm; Techniplast, UK) with temperature and humidity control (21 ±1.5 °C, 50% ±8%) and an alternating 12h light dark cycle (lights on at 0700). Rats had *ad libitum* access to food (Teklad Global 18% protein diet, Harlan, UK) throughout the study. Access to water was restricted during the CER procedure (see details below), but was available *ad libitum* during all other stages of the study. All rats were habituated to handling by experimenters for at least 5 days prior to any experimental procedure. All experimental procedures were conducted during the light phase and in accordance with the requirements of the United Kingdom (UK) Animals (Scientific Procedures) Act 1986, approved by the University of Nottingham’s Animal Welfare and Ethical Review Board (AWERB) and run under the authority of Home Office project license 30/3357.

### Stereotaxic implantation of guide cannulae into the VH

Rats were anaesthetised using isoflurane delivered in oxygen (induced with 5% and maintained at 1.5-3%; flow rate 1L/min) and then placed in a stereotaxic frame. A local anaesthetic (EMLA cream, AstraZeneca, UK) was applied to the ear bars to minimise discomfort. A gel was used (Lubrithal; Dechra, UK) to prevent the eyes from drying out during surgery. After incision of the scalp, bilateral infusion guide cannula (stainless steel, 26 gauge, 8.5mm below pedestal, Plastics One, USA) were implanted through small pre-drilled holes in the skull. The stereotaxic coordinates for the injections were 5.2mm posterior, ±4.8mm lateral from the midline and 6.5mm ventral from the dura for infusions into the VH, as in McGarrity et al. (2017). Stainless steel stylets (33 gauge, Plastics One, USA), complete with dust cap, were placed into the guide cannula and protruded 0.5mm beyond the tips of the guide cannula to prevent occlusion. Dental acrylic (flowable composite; Henry Schein, Germany) and four stainless steel screws were used to fix the guide cannulae to the skull. The scalp incision was stitched around the acrylic pedestal to reduce the open wound to a minimum. All rats were injected with perioperative analgesia (Rimadyl, Large Animal Solution, Zoetis, UK; 1:9 dilution; 0.1ml/100g s.c). At the end of surgery, rats were injected with 1ml of saline (i.p) to prevent dehydration. Antibiotics were administered on the day of surgery and subsequently every 24h for the duration of the study (Synulox; 140mg amoxicillin, 35mg clavulanic acid/ml; 0.02ml/100g s.c, Pfizer, UK). After surgery, rats were allowed at least 5 days of recovery before any further experimental procedures were carried out. During this period, rats underwent daily health checks and were habituated to the manual restraint necessary for drug microinfusions.

### Microinfusions into the VH

Rats were manually restrained throughout the infusion process. Stylets were replaced with infusion injectors (stainless steel, 33 gauge, Plastics One, USA), which extended 0.5 mm below the guide cannula tips into the VH. Injectors were connected via flexible polyethylene tubing to 5-µl SGE micro-syringes mounted on a microinfusion pump (sp200IZ, World Precision Instruments, UK). A volume of 0.5µl/side of either 0.9 % sterile saline (vehicle) or picrotoxin (150ng/0.5µl/side; Sigma Aldrich, UK) in saline was infused bilaterally over the course of 1min, as in our previous studies (McGarrity et al., 2017). The movement of an air bubble, which was included in the tubing, was monitored to ensure the solution had been successfully injected into the brain. Injectors were removed and replaced by the stylets 60s after the end of infusion to allow for tissue absorption of the infusion bolus. The timing of infusions in relation to behavioural testing is described below, in the Experimental design section.

In our previous work, the dose of picrotoxin (150ng/0.5µl/side) used did not cause seizure-related behavioural signs or electrophysiological signs of hippocampal seizures in local field potential recordings in anaesthetised rats (McGarrity et al., 2017). However, picrotoxin has the potential to cause epileptiform activity in the hippocampus (Qaddoumi et al., 2014). Therefore, all rats receiving infusions were monitored carefully during and after infusion for behavioural signs potentially related to seizure development, including facial twitching, wet-dog shakes, clonic limb movement, motor convulsions and wild jumping (Luttjohann et al., 2009; Racine, 1972).

### CER procedure with a pre-exposure phase to measure aversive conditioning and its latent inhibition

We used a CER procedure described by Nelson et al. (2011). The procedure, which will be described in detail below, involved water deprivation, shaping (1 d) and pre-training of the rats to drink from spouts in conditioning chambers (5 d), followed by pre-exposure to a light (the prospective CS) in conditioning chambers (or exposure to the conditioning chamber without CS pre-exposure in the non-pre-exposed comparison group) (1 d), conditioning during which the CS was paired with an electric footshock, reshaping (1 d) to re-establish drinking after conditioning and testing (1 d) of the lick suppression induced by CS presentation following conditioning (for an outline of the CER stages, also see Figs 2A and 3A). Suppression of licking for water by the CS was used to measure the CER. LI is reflected by a reduced CER, i.e. less suppression of licking for water, in the pre-exposed (PE) as compared to the non-pre-exposed (NPE) group.

**Figure 1.**
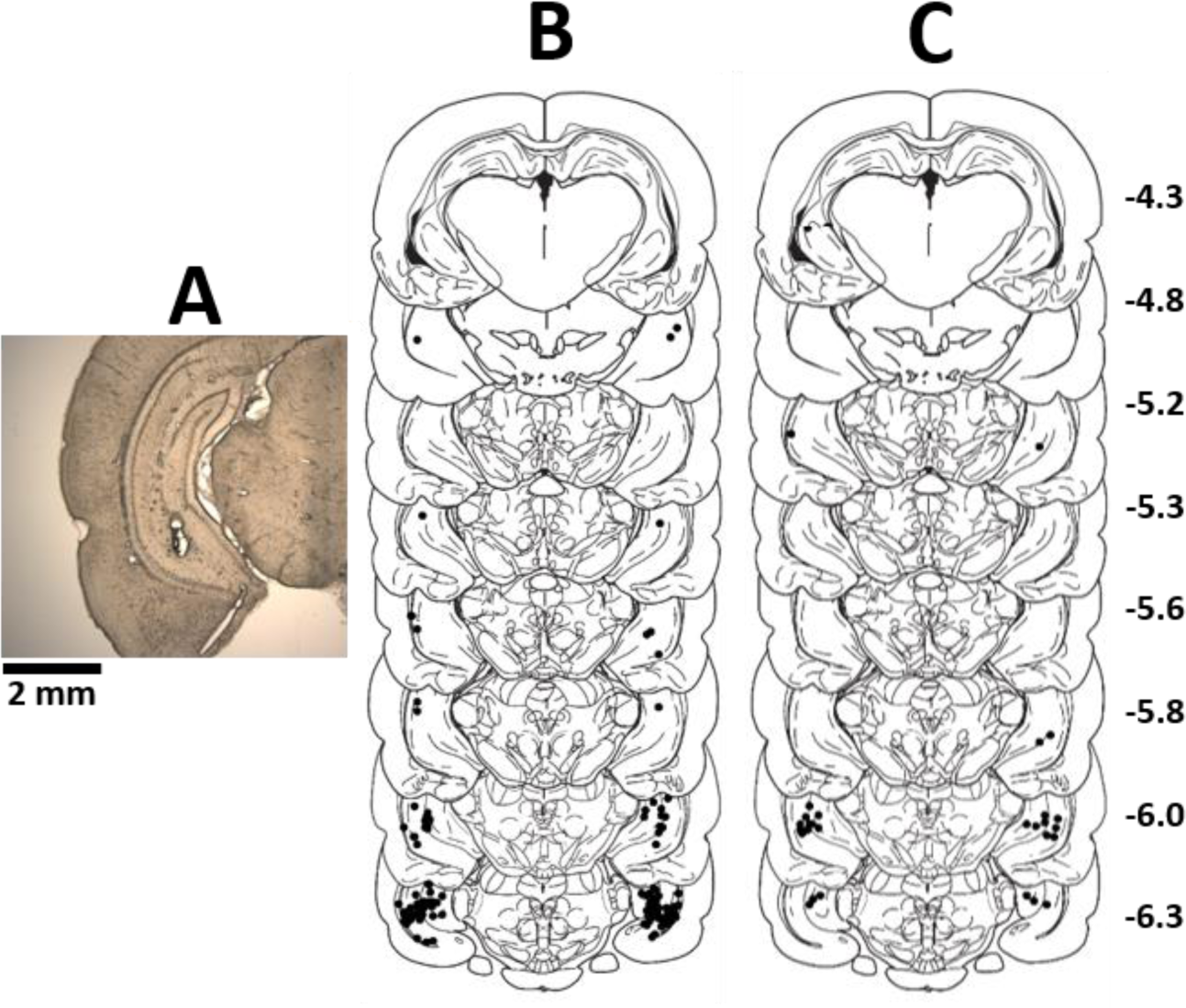
Infusion sites in the VH. **A:** Illustrative coronal brain section showing infusion sites in the VH. Approximate locations of infusion cannula tips (black dots) mapped onto coronal sections adapted from the Paxinos and Watson (1998) rat brain atlas for rats in Experiment 1 **(B)** and 2 **(C)**. Numbers on the right indicate posterior distance from bregma in mm.

**Figure 2.**
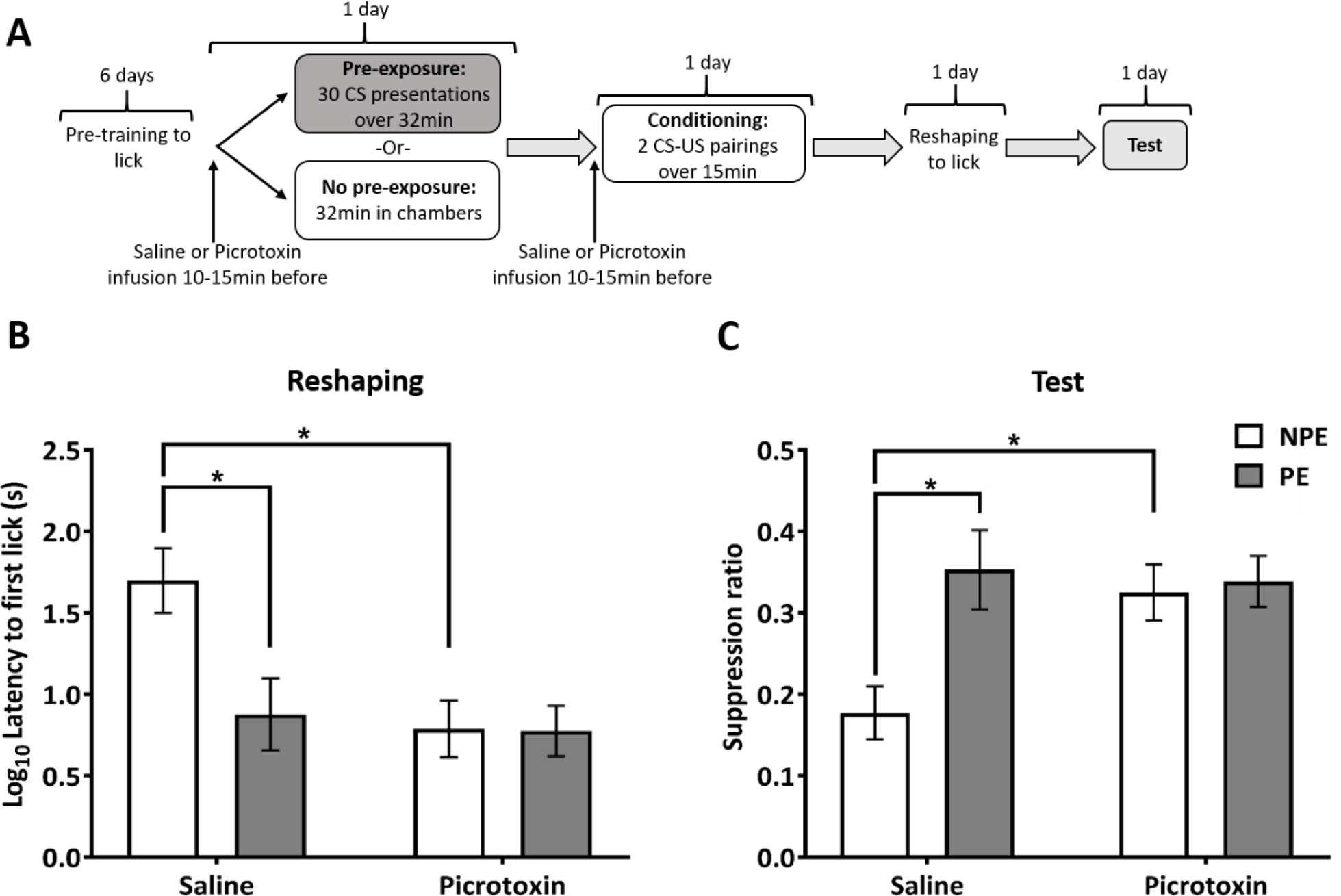
Experiment 1: Ventral hippocampal disinhibition during pre-exposure and conditioning impairs the acquisition of contextual and elemental fear conditioning. **A:** Design of Experiment 1. **B:** Mean (± SEM) latency to first lick values (s) (log transformed) in the conditioning chamber following the aversive conditioning session for non-pre-exposed (NPE, white bars) and pre-exposed (PE, grey bars) rats in the saline and picrotoxin groups. Saline NPE rats show longer latencies compared to all other groups indicating increased conditioning to the conditioning context. Picrotoxin-infused rats show reduced latencies compared to saline-infused animals indicating impaired conditioning to the conditioning context. **C:** Mean suppression ratio (±SEM) to the light conditioned stimulus for NPE (white) and PE (grey) rats in the saline and picrotoxin groups. Saline-infused rats displayed LI, with PE rats showing markedly less fear than NPE rats. Picrotoxin-infused rats show similarly low levels of fear conditioning in both NPE and PE groups reflecting picrotoxin infusion abolished conditioning to the CS. * Asterisks indicate statistically significant differences between groups (*F* > 9, *p* < 0.005; simple main effects analysis following significant interaction of infusion and pre-exposure).

**Figure 3.**
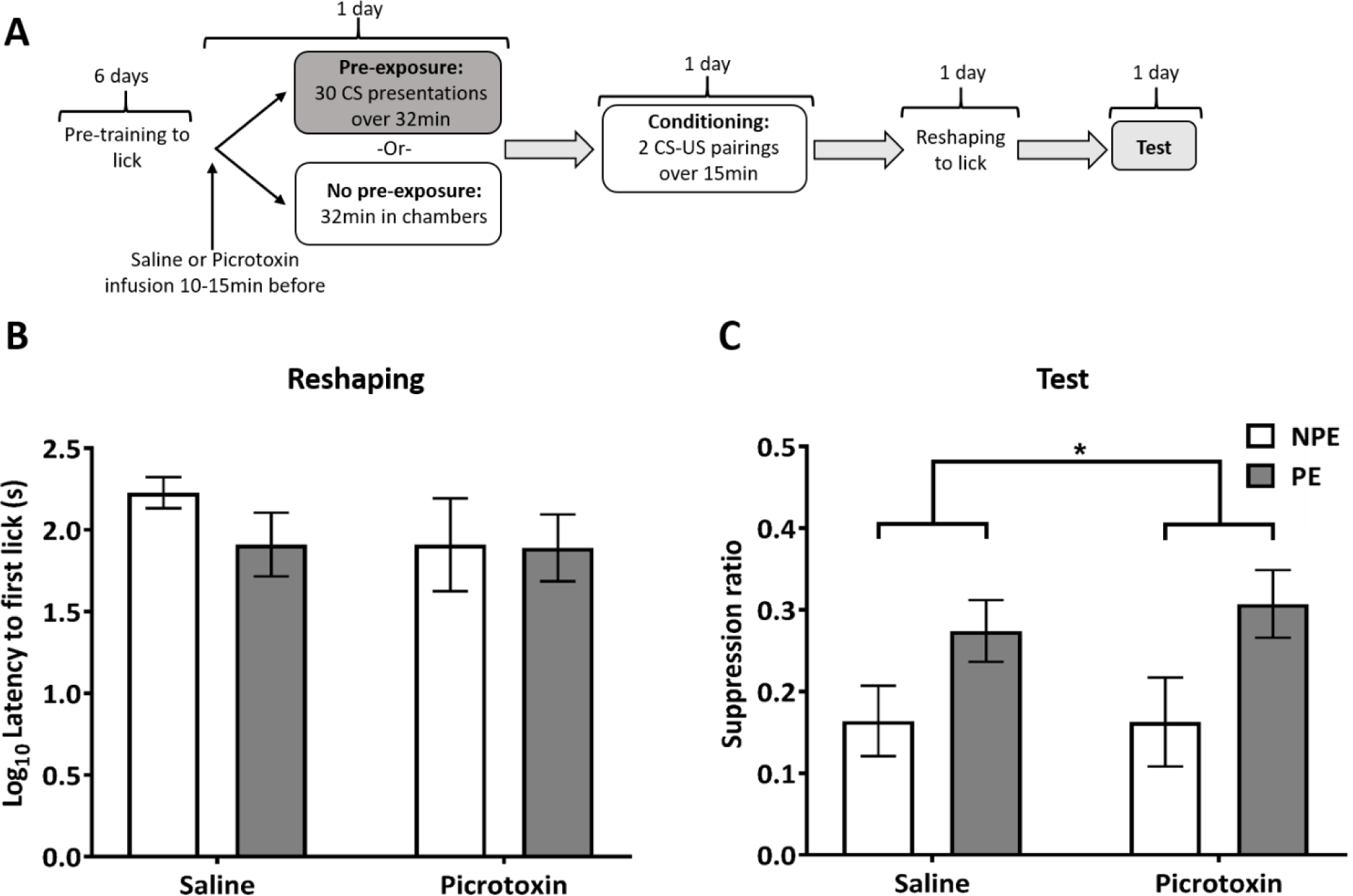
Experiment 2: VH disinhibition during pre-exposure does not impair the acquisition of LI. **A:** Design of Experiment 2, with the time point of the VH picrotoxin or saline infusion before the pre-exposure stage indicated. **B:** Mean (± SEM) latency to first lick (s) (log transformed) in the conditioning chamber, during reshaping, following the aversive conditioning session for non-pre-exposed (NPE, white bars) and pre-exposed (PE, grey bars) rats in the saline and picrotoxin groups. All groups show similar levels of contextual conditioning, indicated by similar latencies to first lick. **C:** Mean suppression ratio (±SEM) to the light conditioned stimulus (CS) for control NPE (white) and PE (grey) rats in the saline and picrotoxin groups. Pre-exposure reduced fear responding to the CS in both saline and picrotoxin-infused rats compared to NPE rats, reflecting LI in both saline and picrotoxin-infused rats. * Asterisk indicates significant main effect of pre-exposure in saline-infused and picrotoxin-infused rats (*F* > 8, *p* < 0.01).

#### Apparatus

Four identical fully automated conditioning chambers including sound attenuating cases and ventilation fans (Cambridge Cognition, UK) were used. The inner chambers consisted of a plain steel box (25cm x 25cm x 22cm) with a Plexiglas door (27cm x 21cm). The floor of the inner conditioning chamber comprised of a shock delivery system, consisting of 1cm spaced steel bars. These were positioned 1cm above the lip of a 7cm deep sawdust tray. Mounted 5cm above the grid floor was a waterspout connected to a lickometer supplied by a water pump. Licks were registered by breaking a photo beam within the spout, which triggered water delivery of 0.05ml per lick. The spout was only illuminated when water was available. Three wall mounted lights and the house light flashing on (0.5s) and off (0.5s) for 5s functioned as the CS. Scrambled foot-shock of 1mA intensity for 1s provided the unconditioned stimulus (US). The shock was delivered through the grid floor by a constant current shock generator (pulsed voltage: output square wave 10ms on, 80ms off, 370V peak under no load conditions; MISAC Systems, UK). Stimulus control and data collection were recorded using an Acorn RISC computer programmed in basic with Arachnid extension (Cambridge Cognition, UK).

#### Behavioural procedure

##### Water restriction

One day prior to behavioural testing, rats were water restricted for between 18-22h. Subsequently, they received 1h and 15min of *ad libitum* access to water in their home cages for the duration of the experiment, once daily testing was completed and in addition to access to water in the conditioning chambers.

##### Shaping and pre-training

Rats were shaped for 1 day until all rats drank from the waterspout and were assigned an individual conditioning chamber for the whole CER procedure. Subsequently, rats were given a 15min session (timed from first lick) per day for 5 days to drink from the waterspout. During the sessions, the waterspout was illuminated throughout, but no other stimuli were present. Total number of licks was recorded during each session to assess any pre-existing differences in drinking prior to infusions.

##### Pre-exposure

The PE rats received 30 5s flashing light CS presentations with an average inter-stimulus interval of 60s (32min session duration). The NPE control rats were confined to the conditioning chamber for an identical period of time without receiving any CS presentations. Water was not available during the session and the waterspout was not illuminated.

##### Conditioning

One day after pre-exposure, rats were conditioned by two light-foot shock pairings, with the foot shock (1mA/1s) delivered immediately following the termination of the flashing light (5s). The first light-shock pairing was presented after 5min had elapsed and the second pairing 5min after the first, followed by a further 5min in the chamber, resulting in an overall session duration of 15min. Water was not available during the session and the waterspout was not illuminated for the duration of the session.

##### Reshaping

The day after conditioning, rats were reshaped using the same procedure as used during the initial shaping. This was to re-establish drinking behaviour after the conditioning session. Latency to first lick during reshaping was used as a measure of contextual fear conditioning to the chamber (Nelson et al., 2011; Nelson et al., 2013).

##### Test

The day after reshaping, rats underwent a test session to assess conditioning to the CS. During the test session, water was available throughout and the waterspout was illuminated. Once the rats had performed 50 licks, the CS was presented continuously for 15min. The time taken to complete 50 licks before CS presentation (excluding latency to first lick) provides a measure of individual baseline variation (A period). This time was compared to the time taken to complete 50 licks during CS presentation (B period). A suppression ratio (A / (A+B)) was used to assess the overall level of conditioning to the CS, adjusted to individual variation in drinking, where a higher ratio represents a low level of fear conditioning (with a value of 0.5 or higher indicating no conditioning at all) and a ratio closer to 0 represents a high level of conditioning to the CS (Nelson et al., 2011; Nelson et al., 2012).

### Verification of cannula placements

After behavioural experiments, rats were deeply anaesthetised with sodium pentobarbital (Dolethal, Vetoquinol, UK) and were transcardially perfused with 0.9% saline followed by 4% paraformaldehyde in saline. Subsequently brains were removed and stored in 4% paraformaldehyde. Brains were sliced at 80µm thickness using a vibratome and placed on microscope slides. Injector placements were identified using light microscopy and mapped onto coronal sections of a rat brain atlas (Paxinos & Watson, 1998).

### Experimental design

Both Experiments 1 and 2 were run in a between subjects design with a target sample size for both experiments of 16-18 per group. This sample size would give a power of > 80% to detect effect sizes of Cohen’s *d*=1 for differences between groups (using between-subjects pairwise comparisons, two-tailed, with a significance threshold of *p*<0.05; G*Power (Faul et al., 2007)), which has been suggested to be appropriate for neurobiological studies of aversive conditioning (Carneiro et al., 2018). Experiment 1 was run in 3 identical series, each including 24 rats. Experiment 2 was planned to comprise of 2 series, each containing 32 rats, but was ended after the first series. The second series was unnecessary, as there was clearly no evidence that the target effect size the study was powered for could be achieved (Neumann et al., 2017).

Rats were allocated to experimental groups according to a randomised block design. Two of the four rats in each cage were randomly assigned to the saline and the other two to the picrotoxin infusion group, and subsequently one rat of each pair was randomly assigned to either PE or NPE groups. The experimenters were blinded with respect to the infusion group allocation at the start of the experiment. In both experiments, several rats had to be excluded from the analysis of the whole experiment or some later stages of the experiment. During Experiment 1, 13 rats fell ill, with presumed meningitis, before reshaping, whilst a further 2 rats fell ill after reshaping and prior to the test session; two additional rats had blocked guide cannulae after surgery and before behavioural testing, resulting in exclusion from the experiment; another rat showed extended convulsive seizures after picrotoxin infusion prior to conditioning. During Experiment 2, one rat died during surgery and a further three rats fell ill, with presumed meningitis, prior to the reshaping session. The final sample sizes contributing to the analysis of performance measures at the different test stages in Experiment 1 and 2 are shown in Table 1.

**Table 1.**
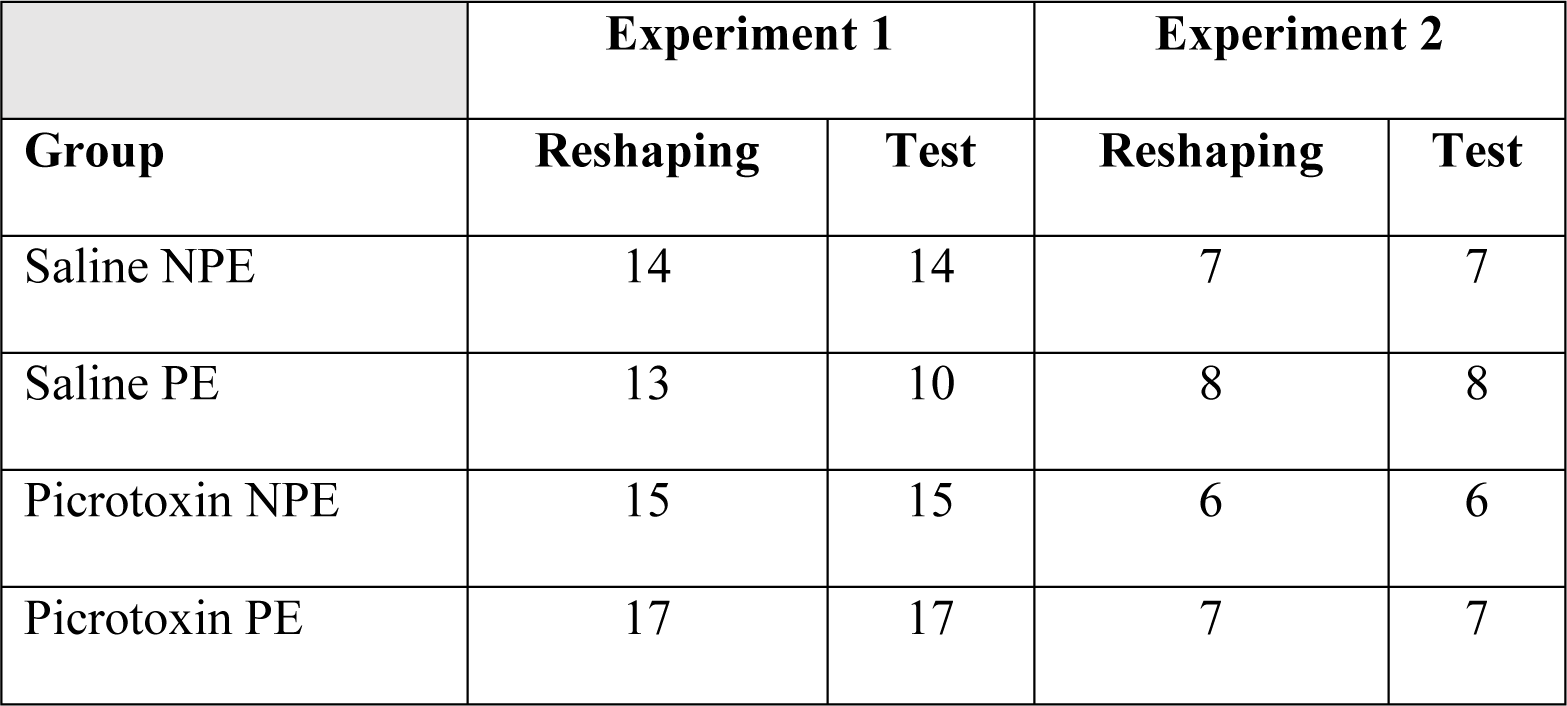
Final number of rats included in data analysis per group for each stage of both experiments.

|  | **Experiment 1** |  | **Experiment 2** |  |
| --- | --- | --- | --- | --- |
| **Group** | **Reshaping** | **Test** | **Reshaping** | **Test** |
| Saline NPE | 14 | 14 | 7 | 7 |
| Saline PE | 13 | 10 | 8 | 8 |
| Picrotoxin NPE | 15 | 15 | 6 | 6 |
| Picrotoxin PE | 17 | 17 | 7 | 7 |

In Experiment 1, VH drug infusions took place before both pre-exposure and conditioning sessions (Fig. 2A), whereas, in Experiment 2, drug infusions took place before pre-exposure only (Fig. 3A). Rats were infused in batches of two pairs, by two experimenters, with each pair including one rat to receive saline and one rat to receive picrotoxin infusions. The two experimenters infused one pair, then the second pair, and testing started 10min after the infusions for both rats of the second pair had been completed. This meant that all rats had a 10-15 min period between the end of the infusion and the start of behavioural testing. The timing of behavioural procedures after intracerebral infusions was based on our previous electrophysiological measurements to capture the peak effect of hippocampal picrotoxin on neuronal firing following infusion (McGarrity et al., 2017).

### Statistical analysis

The measures taken during the CER experiments were analysed using a 2×2 analysis of variance (ANOVA) with between subject factors of pre-exposure group (NPE/PE) and drug infusion (Saline/Picrotoxin). All statistical tests and graphs were completed using SPSS (version 23), JASP (JASP Team: version 0.12.2, 2020) and Graphpad prism (version 7) software. The accepted level of significance was *p* < 0.05. Raw latency data (time to first lick during reshaping) or time ‘A’ data (time to 50 licks during test) were log transformed, as they showed unequal variance (Levene’s test, all *F* > 5, *p* < 0.002), to ensure a normal distribution and suitability for parametric analysis (Nelson et al., 2011; Nelson et al., 2012).

## RESULTS

### Cannula placements in the VH

In both experiments, all cannula tips were located within the VH, in coronal brain sections corresponding to between 4.3 and 6.3mm posterior to bregma in the rat brain atlas by Paxinos and Watson (1998) (Fig. 1).

### Experiment 1: VH disinhibition during pre-exposure and conditioning disrupts aversive conditioning

#### Pre-training

Analysis of latencies to lick at the end of pre-training, prior to pre-exposure, showed no overall effect of prospective infusion or pre-exposure group, nor an interaction of these factors (all *F* < 1) (data not shown).

#### Reshaping

VH picrotoxin, compared to saline, infusion reduced latencies to first lick after reintroduction to the conditioning context during the reshaping session in the NPE group, which reflects reduced contextual fear conditioning. In the PE group both saline and picrotoxin groups showed similarly low levels of contextual conditioning, as measured by low latencies to lick, which indicates that pre-exposure to the light CS reduced contextual conditioning in the saline group (Fig. 2B). These observations were supported by a significant infusion x pre-exposure interaction (*F* = 4.7, *p* = 0.034). Further examination of the interaction by simple main effects analysis showed that hippocampal picrotoxin, compared to saline, reduced conditioning in the NPE group (*F* = 11.9, *p* = 0.001), but this was not apparent in the PE group, due to a floor effect where both saline and picrotoxin rats showed similarly low conditioning (*F* < 1). In addition, pre-exposure to the CS reduced context conditioning in saline-infused rats, reflected by reduced latencies in the PE group as compared to the NPE group (*F* = 9.0, *p* = 0.004). This effect was not present in picrotoxin-infused rats (*F* < 1), probably reflecting a floor effect, i.e. the already low latencies in the picrotoxin rats.

#### Test

There was no difference in time to 50 licks before CS presentation (Time A) between infusion groups and pre-exposure groups (any effect or interaction involving infusion or pre-exposure: all *F* < 3, *p* > 0.09; data not shown). The group differences in latency to first lick that were evident at reshaping were not present during the test stage, probably reflecting extinguished contextual conditioning in the saline NPE group. The suppression ratios during the light test revealed that hippocampal disinhibition markedly disrupted conditioning to the CS in the NPE group, but did not affect conditioning in the PE group, i.e. there was no evidence that hippocampal disinhibition had affected LI (Fig. 2C). In saline-infused rats, the suppression ratio was markedly increased in the PE compared to the NPE group, reflecting reduced conditioning, i.e. LI (Fig. 2C, left). This difference between PE and NPE groups was not apparent in the picrotoxin-infused rats (Fig. 2C, right). However, this was due to picrotoxin-infused NPE rats showing a markedly higher suppression ratio than saline-infused NPE rats, reduced conditioning to the light CS (compare white bars in Fig. 2C). In contrast, suppression ratios were similar in picrotoxin and saline-infused PE rats (compare grey bars in Fig. 2C). Thus there was no evidence that hippocampal disinhibition reduced the impact of CS pre-exposure on conditioning. These observations were supported by a significant infusion x pre-exposure interaction (*F* = 4.142, *p* = 0.047). Further examination of the interaction by simple main effects analysis revealed a main effect of infusion in the NPE group (*F* = 10.014, *p* = 0.003) reflecting increased suppression ratio, i.e. reduced conditioning, caused by picrotoxin, compared to saline, whereas there was no effect of infusion in the PE group (*F* < 1). This resulted in the absence of a difference between PE and NPE in the picrotoxin-infused rats (*F* < 1), whereas saline-infused rats showed markedly higher suppression in the PE compared to the NPE group (*F* = 12.111, *p* = 0.001).

In Experiment 1, the VH was disinhibited during pre-exposure and conditioning, but not during reshaping and test. Therefore, the impaired fear conditioning evident during reshaping and test sessions could reflect state dependence, i.e. that information learned in one neural state can, in some cases, only be retrieved/expressed in the same state (Overton, 1964). To rule this out would require showing that fear expression is disrupted if the VH is disinhibited both during conditioning and the test expression of fear, but the interpretation of this finding would be difficult because the VH has been implicated in the expression of conditioned fear (Sierra-Mercado et al., 2011). However, several studies have shown that state dependent learning does not account for the conditioning deficits caused by local drug microinfusions into specific brain sites, including the mPFC, amygdala, and dorsal hippocampus (Bast et al., 2003; Guarraci et al., 2000; Pezze et al., 2003). In addition, previous experiments using a similar 3-stage fear conditioning paradigm to study systemic drug effects on LI found no evidence for state-dependent effects (Barad et al., 2004). Altogether, a specific impairment in neural mechanisms underlying the formation of fear memory is the most plausible account for the reduced conditioned suppression following VH disinhibition during conditioning.

### Experiment 2: VH disinhibition during pre-exposure alone does not affect conditioning or LI

#### Pre-training

Analysis of latencies to lick at the end of pre-training, prior to pre-exposure, showed no overall effect of prospective infusion (*F* =2.9, *p* = 0.104) or pre-exposure group (*F* < 1), and there was no interaction of these factors (*F* < 1) (data not shown).

#### Reshaping

Hippocampal picrotoxin infusion only at pre-exposure had no effect on conditioning to the context, as reflected by latencies to first lick during reshaping, and there was no difference between pre-exposure groups (all main effects and interactions, *F* < 1.5, *p >* 0.2) (Fig. 3B). The latter contrasts with the finding in Experiment 1, that pre-exposure reduced latencies to first lick in saline-infused rats (Fig. 2B).

#### Test

There were no differences in the A period (time to 50 licks before CS presentation) between infusion and pre-exposure groups (all main effects and interactions, *F* <1.2, *p* > 0.30) (data not shown). Both drug infusion groups showed similar fear conditioning to the light CS, reflected by similar suppression ratios, and robust LI, reflected by higher suppression ratios in the PE compared to the NPE groups (Fig. 3C). This was supported by an effect of pre-exposure group (*F* = 8.44, *p* = 0.0078), without a main effect or interaction involving infusion group (both *F* < 1).

### Seizure-related behavioural effects of hippocampal picrotoxin

In several rats receiving hippocampal picrotoxin infusions in Experiment 1 (20 out of 32 rats receiving picrotoxin) and Experiment 2 (6 out of 15 rats receiving picrotoxin), we observed seizure-related behavioural signs, including facial twitching, wet-dog shakes and wild running, which can often be observed before full motor seizures (Luttjohann et al., 2009; Racine, 1972). These effects were observed within 5 min after the end of the picrotoxin infusion. They typically subsided within 30-45 min, after which rats showed no further adverse effects, with the exception of one rat, which showed continued uncontrollable clonic limb movement and was culled. We never observed these signs following saline infusions. Table 2 shows how many rats showed any of these seizure-related effects after the two picrotoxin infusions of Experiment 1 or the one picrotoxin infusion of Experiment 2. Although GABA network dysfunction, including in the hippocampus, is strongly implicated in the onset of seizures (Avoli & de Curtis, 2011) and the VH is a particularly seizure prone brain region, showing the earliest seizure activity in the pilocarpine rat model of seizures (Toyoda et al., 2013), our previous studies using the same dose of picrotoxin as in the present study did not reveal seizure-related effects in Lister Hooded (McGarrity et al., 2017) or Wistar rats (Bast et al., 2001a). Given that stress substantially facilitates hippocampal seizures (Joels, 2009; Manouze et al., 2019), the seizure-related effects of hippocampal picrotoxin infusions in the present study may reflect that, in contrast to our previous studies involving hippocampal picrotoxin infusions, rats in the present study were exposed to water restriction and foot shocks as part of the CER procedure.

**Table 2.** Seizure-related behavioural signs observed after VH picrotoxin microinfusions. The type of behaviour observed is indicated in column one. Total number of rats experiencing seizure-related behaviour signs overall during Experiment 1 or 2 is shown in column two. The number of rats experiencing seizure-related signs during Experiment 1 is detailed in column three, with these signs separated to show the effects after the two individual infusions in columns 4 and 5. Column 6 details the total number of rats showing seizure-related signs after the one infusion of Experiment 2.

| Observed behaviour | Overall<br>total | Experiment<br>1 total | Experiment 1 |  | Experiment 2<br>total |
| --- | --- | --- | --- | --- | --- |
|  |  |  | Infusion 1 | Infusion 2 |  |
| Facial twitching | 3 | 3 | 1 | 2 | 0 |
| Wet dog shakes | 19 | 15 | 11 | 7 | 4 |
| Wild running | 10 | 9 | 8 | 2 | 1 |
| Clonic limb movement | 1 | 0 | 0 | 0 | 1 |

Importantly, additional analyses excluding rats that showed seizure-related behaviours during conditioning still revealed a disruption of contextual and elemental fear conditioning in rats with VH disinhibition compared to saline-infused control rats in Experiment 1. More specifically, in NPE rats, VH disinhibition still reduced latencies to lick during reshaping (strong trend towards interaction infusion X pre-exposure: *F* = 3.614, *p* = 0.0636; simple main effect of infusion in NPE group: *F* = 5.330, *p* = 0.026) and reduced conditioned suppression during test (interaction infusion X pre-exposure: *F* = 4.933, *p* = 0.0317; simple main effect of infusion in NPE group: *F* = 8.310, *p* = 0.006). Therefore, the disruption of fear conditioning by hippocampal disinhibition was not a consequence of seizure-related behavioural effects during conditioning.

## DISCUSSION

In Experiment 1, VH disinhibition by picrotoxin during pre-exposure and conditioning markedly reduced fear conditioning to the CS and, therefore, any reduction of fear conditioning in the PE compared to NPE group, which would indicate LI, could not be detected. Picrotoxin and saline-infused rats in the PE group did not differ, showing similarly low conditioning, which does not support the hypothesis that hippocampal disinhibition affected salience modulation. In addition to disrupting conditioning to the CS, VH disinhibition also impaired contextual fear conditioning. In Experiment 2, which specifically examined the impact of hippocampal disinhibition during pre-exposure alone, there was no evidence for any impact on LI.

### Pre-exposure-induced reduction of contextual fear conditioning

In Experiment 1, the saline-infused PE rats showed shorter latencies to the first lick than NPE rats, reflecting reduced fear conditioning to the context. This could reflect that the novelty of the light stimulus enhanced memory formation (Duszkiewicz et al., 2019; King & Williams, 2009; Lisman & Grace, 2005) in the NPE group. The reduced context conditioning in PE compared to NPE saline-infused rats was not evident in Experiment 2. This could be accounted for by a ceiling effect, i.e. higher levels of context conditioning, in Experiment 2, which may have masked any further novelty-induced enhancement of context conditioning in the NPE group. In previous studies, un-operated rats showed stronger fear conditioning than cannulated rats that received hippocampal saline infusions, in terms of conditioned freezing (Zhang et al., 2001) and lick suppression (Zhang et al., 2000), suggesting that the infusion procedure itself might reduce fear conditioning. Therefore, the stronger conditioning in Experiment 2 may partly reflect that, in contrast to Experiment 1, the rats did not receive drug infusions immediately before conditioning.

### VH disinhibition during pre-exposure and conditioning markedly reduces fear conditioning without affecting LI

In Experiment 1, VH disinhibition during both pre-exposure and conditioning markedly reduced fear conditioning to the CS in the NPE group, resulting in similarly low levels of conditioning in both the NPE and PE groups. Whilst there was no evidence for LI following VH disinhibition, the absence of LI was not due to increased conditioning in the PE group, which would reflect aberrant salience allocation, but instead was due to reduced conditioning in the NPE group. Similar to the present study, Pouzet et al. (2004), using a comparable LI paradigm, demonstrated VH NMDA stimulation reduced conditioned suppression in the NPE group, although there was also some evidence for disrupted LI with a trend towards greater conditioned suppression in PE compared to NPE rats. Moreover, studies in the MAM rat model of schizophrenia, which shows a loss of parvalbumin GABA interneurons and hyperactivity in the VH, also reported the absence of LI, which was mediated by reduced conditioning in the NPE group (Flagstad et al., 2005; Lodge et al., 2009).

### Disruption of elemental and contextual fear conditioning by VH disinhibition might reflect disruption of regional and distal processing

The impairments in fear conditioning to the CS and the context by VH disinhibition are likely mediated at the conditioning stage, which is supported by the finding in Experiment 2 that disinhibition during pre-exposure alone did not affect conditioning. Impaired fear conditioning may reflect disrupted processing within the VH itself and in connected sites (Bast et al., 2017). Lesions, temporary inactivation by the sodium channel blocker TTX, and NMDA stimulation of the VH have been found to disrupt both contextual and elemental fear conditioning (Bast et al., 2001b; Czerniawski et al., 2012; Kjelstrup et al., 2002; Maren, 1999; Zhang et al., 2001). However, functional inhibition of the VH by the GABA agonist muscimol only disrupts contextual, but not elemental, conditioning, suggesting that neurons within the VH are mainly required for contextual fear conditioning (Bast et al., 2001b; Zhang et al., 2014). Therefore, the impaired contextual fear conditioning in the present study may reflect that disinhibition disrupts VH processing, whereas disrupted elemental fear conditioning is consistent with the idea that regional disinhibition can disrupt processing in VH projection sites (Bast et al., 2017), which have been implicated in elemental fear conditioning (see next paragraph). However, changes in dorsal hippocampal function, which is necessary for contextual fear conditioning and has been suggested to produce the underlying contextual representation (Anagnostaras et al., 2001; Bast et al., 2003; Hunsaker & Kesner, 2008; Matus-Amat et al., 2004), may also contribute to contextual fear conditioning deficits caused by VH disinhibition. VH disinhibition might disrupt dorsal hippocampal function by way of intra-hippocampal inhibitory longitudinal connections (Sik et al., 1994; Sik et al., 1997). In line with this suggestion, our recent metabolic imaging study showed that VH disinhibition activated the VH, but deactivated the dorsal hippocampus (Williams et al., 2019).

The VH also sends strong projections to the amygdala, mPFC and septum (Cenquizca & Swanson, 2007; Hoover & Vertes, 2007; Pitkanen et al., 2000; Risold & Swanson, 1997), all of which are components of a brain circuit controlling conditioned fear responses to elemental stimuli (Tovote et al., 2015). The amygdala is a key component of the fear conditioning circuit and is thought to play a crucial role in the CS-US association and in conveying conditioned fear information to downstream effector sites (Duvarci & Pare, 2014; LeDoux, 2000). Thus, VH disinhibition, by causing aberrant drive of projections to the amygdala, could disrupt the processing of CS-US associations underlying conditioned fear. The mPFC is mainly thought to be required for the expression of cue conditioning and not its acquisition (Corcoran & Quirk, 2007; Morgan et al., 1993; Pezze et al., 2003), although inactivation of the rostral anterior cingulate cortex disrupted the acquisition of cue fear conditioning (Bissiere et al., 2008). The anterior cingulate cortex does not receive direct VH projections (Bian et al., 2019; Jay & Witter, 1991), but aberrant drive of VH projections to the mPFC might contribute to the disruption of elemental fear conditioning by way of regional connectivity within the mPFC (Jones et al., 2005). The lateral septum receives strong glutamatergic VH projections (Cenquizca & Swanson, 2007; Risold & Swanson, 1997) and is required for the acquisition of elemental fear conditioning (Calandreau et al., 2007). Additionally, hippocampo-lateral septum neurotransmission has been implicated in the modulation of the strength of CS-US associations and adaptive acquisition of conditioned fear responses (Calandreau et al., 2010; Desmedt et al., 2003). Our recent neuroimaging study showed that VH disinhibition caused significant neural activation changes in the amygdala, mPFC and LS (Williams et al., 2019) and, therefore, VH disinhibition could disrupt elemental fear conditioning by disrupting information processing at these projection sites.

### Hippocampal disinhibition during pre-exposure has no effect on the formation of LI

While aberrant dopamine transmission is thought to disrupt LI by interfering with the effect of pre-exposure during conditioning (Morrens et al., 2020; Young et al., 2005), stimulation and inhibition of GABA receptors disrupted LI formation at the pre-exposure stage (Feldon & Weiner, 1989; Lacroix et al., 2000). However, the lack of effect on LI acquisition by VH disinhibition during pre-exposure in Experiment 2 suggests that sites outside the VH mediate the disruption of LI formation by systemic GABA receptor blockade during pre-exposure (Lacroix et al., 2000). Moreover, although VH disinhibition caused aberrant mPFC activation (Williams et al., 2019) and deficits in mPFC-dependent attention (McGarrity et al., 2017), our present findings show that VH disinhibition does not affect mPFC-dependent processing involved in LI formation during pre-exposure (Lingawi et al., 2017; Lingawi et al., 2018). In line with this, mPFC disinhibition during pre-exposure and conditioning did not disrupt LI formation (Enomoto et al., 2011; Piantadosi & Floresco, 2014). This is consistent with the idea that different prefrontal functions can display distinct relationships to prefrontal neural activity (Bast et al., 2017), with LI formation disrupted only by reductions (Lingawi et al., 2018), but not increases (Enomoto et al., 2011; Piantadosi & Floresco, 2014), in prefrontal activity, whereas sustained attention requires balanced levels of prefrontal activity (Pezze et al., 2014).

Although the present experiments do not support the hypothesis that ventral hippocampal disinhibition during pre-exposure affects LI, deactivation of the ventral subiculum during pre-exposure disrupted LI in a conditioned taste aversion paradigm, demonstrated by increased conditioning in the PE group (Peterschmitt et al., 2005; Peterschmitt et al., 2008). This suggests that LI formation normally requires the ventral subiculum during pre-exposure, but not balanced levels of ventral hippocampal activity.

### Clinical relevance

Our findings do not support the hypothesis that VH disinhibition contributes to LI impairments in schizophrenia. Apart from impairments in LI and other aspects of salience modulation (Roiser et al., 2009; Roiser et al., 2013), fear conditioning deficits have been reported in schizophrenia (Holt et al., 2009; Holt et al., 2012). Such deficits were suggested to contribute to difficulties in differentiating relevant from irrelevant stimuli (Hofer et al., 2001; Jensen et al., 2008) and were associated with negative symptoms (Holt et al., 2012). Previous findings have implicated prefrontal disinhibition in aversive conditioning deficits in schizophrenia (Piantadosi & Floresco, 2014). Our findings suggest that hippocampal disinhibition also contributes to deficits in aversive conditioning.

## Acknowledgements

This work was supported by the Biotechnology and Biological Sciences Research Council (BBSRC) Doctoral Training Programme (DTP) at the University of Nottingham (Standard DTP PhD studentships to SAW and RH; CASE PhD studentships to MG and CT) and the Medical Research Council (MRC) Integrated Midlands Partnership for Biomedical Training (IMPACT) (PhD studentship to JL).

